# Melanomagenesis driven by a kinase-dead mutant BRAF lacking codons 600-618

**DOI:** 10.1101/2021.08.20.457149

**Authors:** Megan A. Girtman, Craig S. Richmond, Meagan E. Campbell, William M. Rehrauer, Paraic A. Kenny

## Abstract

While considerable success has been achieved with treating BRAF-V600E mutant tumors in several organ systems, tumors expressing non-V600E BRAF mutations remain a significant clinical challenge. Such atypical BRAF mutations are classified based on their kinase activity, RAS-dependence and their requirement for dimerization to function. We identified an unusually large in-frame deletion in a melanoma patient, encompassing codons 600-618 in the kinase domain. This patient experienced rapid disease progression when treated with dual BRAF/MEK inhibition. To determine whether and how such a large in-frame BRAF deletion might be activating, we ectopically expressed it in HEK293 cells and monitored BRAF pathway activation and response to BRAF and MEK inhibitors. We demonstrate that this mutation results in a kinase-deficient BRAF mutant which, nevertheless, activates MEK/ERK signaling in a dimerization-dependent manner, placing this mutation among the class III BRAF mutants.

## INTRODUCTION

Constitutive activation of the RAS-RAF-MEK-ERK signaling pathway is common in many different types of cancer. 40-66% of malignant melanomas are caused by somatic mutations in BRAF (Davies et al., 2002; Greaves et al., 2013), a member of the rapidly accelerating fibrosarcoma (RAF) kinase family of mammalian cytosolic serine/threonine kinases (ARAF, BRAF, CRAF). RAF proteins are key intermediates that link RAS GTPases to downstream MEK and ERK kinases which promote proliferation and survival. BRAF mutations are also frequently found in colorectal cancer, ovarian cancer and papillary thyroid carcinomas (Wan et al., 2004). The most common BRAF mutation is the V600E mutation which results in strong activation of the BRAF kinase and high levels of phosphorylated and activated ERK (Wan et al., 2004). Melanoma with BRAF^V600E^ mutation is most effectively treated by combined BRAF and MEK inhibition, such as with dabrafenib and trametinib. (Flaherty et al., 2012).

Recently, studies have proposed that BRAF mutants can be categorized into three classes (Yao et al., 2017). The classification is based on kinase activity, RAS-dependency and requirement for dimerization. Specifically, class I BRAF mutants affect amino acid V600 and function as RAS-independent monomers. Class II BRAF mutations function as RAS-independent activated dimers. Neither class I nor class II mutants require upstream stimulation for activity. Class III BRAF mutations are kinase impaired yet cause increased MAPK signaling. Class III melanoma mutants almost always occur in the context of mutationally activated RAS (whether by direct RAS mutation or NF1 deletion/mutation) and have increased binding to activated RAS, resulting in recruitment of wild-type CRAF and activation of downstream signaling (Heidorn et al., 2010; Yao et al., 2017). Currently, 200 BRAF mutant alleles have been identified in human tumors (Zaman et al., 2019). Of these, 30 distinct mutations of BRAF have been functionally characterized as Class I, II or III mutants (Yao et al., 2015). The ability to classify these mutations may provide the opportunity for more mechanism-focused targeted approaches to treat cancer patients with wider spectrum of BRAF mutations.

In 2017, our statewide molecular tumor board (Burkard et al., 2017) evaluated a melanoma specimen from a male patient in his late fifties with rapidly progressing metastatic disease. A large in-frame deletion in the kinase domain of BRAF encompassing amino acids 600-618 was detected. Small 1-2 codon insertions/deletions have been infrequently noted in the region surrounding V600, however larger in-frame deletions such as D600-605 in a case of papillary thyroid cancer (Barzon et al., 2008) are very rare, and the Δ600-618 variant detected here was far larger than any we had seen documented. Given the known pathway activating effects of smaller deletions in that region (Rogiers et al., 2017), the tumor board at that time recommended treatment with dabrafenib and trametinib. The patient experienced rapid disease progression on this regimen.

Because of the observed lack of response to dabrafenib/trametinib in this case and the uncertainly about whether and how such a large in-frame BRAF deletion might be activating, we attempted to functionally characterize it. Given the substantial disruption to the kinase domain, we hypothesized that this mutation would act as a class III BRAF mutation with impaired kinase activity, leading indirectly to increased MAPK activation. We tested this by evaluating pathway activity in cells ectopically expressing either the mutant kinase or a series of BRAF mutants of known function and evaluated sensitivity to a series of clinically used small molecular inhibitors.

## MATERIALS AND METHODS

### Cell Culture

HEK293 cells were grown in Dulbecco’s Modified Eagle Medium (Thermo Fisher Scientific, Waltham, MA) supplemented with 10% fetal bovine serum (Gemini Bio-products, West Sacramento, CA). HEK293 cells were infected with lentivirus encoding BRAF^V600E^, BRAF^D594A^, BRAF^Δ600-618^. All *BRAF* cDNAs included a FLAG tag on the N-terminus.

### Antibodies

Antibodies against the following targets were used: phospho-p44/42 MAPK (ERK1/2) (#4370), p44/42 MAPK (ERK1/2) (#4695) phospho-MEK1/2 (#3958), Total MEK1/2 (#4694S) all purchased from Cell Signaling Technology (Davers, MA), BRAF (Santa Cruz Biotechnology, Dallas, TX) and FLAG M2 (MilliporeSigma, Burlington, MA). Anti-FLAG M2 magnetic beads (MilliporeSigma, Burlington, MA) were used for immunoprecipitation.

### Inhibitors

Sorafenib, dabrafenib and trametinib were purchased from LC Laboratories (Woburn, MA). Drugs were freshly prepared by dissolving in DMSO (MilliporeSigma, Burlington, MA). Cells were treated with 10 μM of drugs for 4 hours at 37°C.

### Western blot analysis

For western blot analysis, protein was harvested from cells plated at 70-80% confluence. Cells were homogenized in Pierce RIPA lysis buffer (Thermo Fisher Scientific, Waltham, MA) containing protease and phosphatase inhibitors (1/100, Thermo Fischer Scientific, Waltham, MA). Lysates were quantified using a BCA assay (Thermo Fisher Scientific, Waltham, MA) and 35 μg of protein were loaded and size-fractionated by SDS-polyacrylamide gel electrophoresis. (Bio-Rad Laboratories, Hercules, CA). Gels were then transferred to PVDF membranes. After blocking in 5% Blotto (Santa Cruz Biotechnology, Dallas, TX) for 60 minutes at room temperature, membranes were incubated with primary antibodies in dilution buffer at 4°C overnight. The blotted membranes were incubated with HRP-conjugated secondary antibodies (1:1000) at room temperature for 1 hr. Protein expression level was detected and visualized using the Enhanced Chemiluminescence (ECL) detection system (Thermo Fisher Scientific, Waltham, MA).

### BRAF immunoprecipitation and MEK phosphorylation assay

An initial western blot was used to estimate relative expression levels of each FLAG-tagged BRAF construct and subsequent immunoprecipitation amounts were normalized to ensure equal input of each FLAG-BRAF protein. Total cell lysates were incubated overnight at 4°C with anti-FLAG M2 magnetic beads (MilliporeSigma, Burlington, MA) in a final volume of 500 μl of lysis buffer. Following this incubation, the cell lysate was removed, and beads were washed 3 times with 500 μl each, icecold cell lysis buffer followed by 1 wash with 500 μl of ice-cold wash buffer (10 mM Tris pH 7.5, 100 mM NaCl, 0.5% NP-40) and 1 wash with 500 μl of ice-cold reaction buffer (20 mM Tris pH 7.5, 20 mM NaCl, 10 mM MgCl_2_, 1 mM MnCl_2_, 1 mM DTT, 20 μM ATP). Following the final wash, beads were resuspended in 33 μl of reaction buffer containing 500 ng of the phosphorylation target, recombinant MEK1 protein (Santa Cruz Biotechnology, Dallas, TX), and incubated for 30 minutes at 30° C with gentle shaking. The kinase assay was terminated by the addition of 7 μl of SDS sample buffer followed by heating to 95°C for 5 min. The samples were resolved by SDS-PAGE, transferred to Immobilon-P membrane (Millipore Corp., Bedford, MA, USA) and probed with a polyclonal anti-phospho-MEK antibody (S217/221) (Cell Signaling Technologies, Davers, MA). Membranes were stripped and reprobed with anti-FLAG to confirm the levels of FLAG-BRAF protein in each lane.

## RESULTS

### BRAF^Δ600-618^ *elicits increased MEK and ERK activity*

To determine whether BRAF^Δ600-618^ could activate the MEK/ERK pathway, we used lentiviral transduction to express BRAF^Δ600-618^ in HEK293 cells. Lysates from cells transduced with empty vector, BRAF^WT^, BRAF^V600E^, and BRAF^D594A^ served as controls. The most common mutant BRAF, BRAF^V600E^ has been described as a class I BRAF mutant (high kinase activity and functions as a monomer). The BRAF^D594A^ has been previously classified as a class III BRAF mutant (Yao et al., 2017). Class III mutations may either have kinase activity lower than wildtype BRAF or no kinase activity, and drive signaling via heterodimerization with wild-type RAF family members (Wan et al., 2004). In Figure 1, cell lysates were probed for BRAF, FLAG, pMEK and pERK. Although ERK pathway activation was markedly lower than detected in BRAF^WT^ or BRAF^V600E^ expressing cells, cells expressing BRAF^Δ600-618^ had increased phosphorylated ERK and MEK when compared to empty vector controls. This activity level was similar to that elicited by BRAF^D594A^, a kinase-dead BRAF mutation with known oncogenic potential (Heidorn et al., 2010). These results indicate that expression of BRAF^Δ600-618^ elevates MEK/ERK activity to levels comparable to those induced by the oncogenic class III mutant BRAF^D594A^, although much less than the class I mutant BRAF^V600E^.

**Figure 1.**
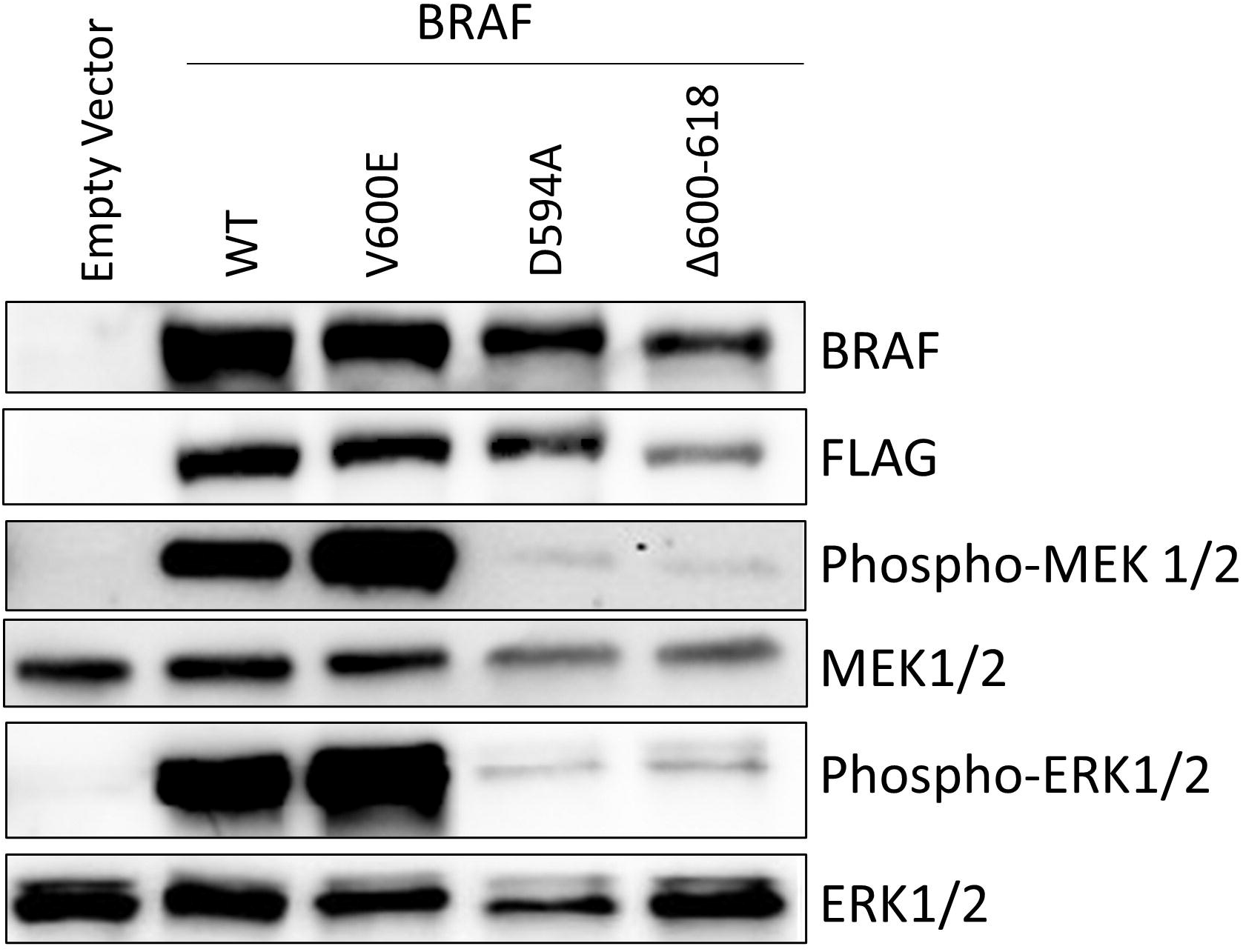
Western blot analysis of MEK/ERK pathway activation following ectopic expression of BRAF^Δ600-618^ and other BRAF mutants in HEK293 cells.

### BRAF^Δ600-618^ *immunoprecipitates have MEK phosphorylating activity*

To determine whether BRAF^Δ600-618^ directly activates MEK by phosphorylation or whether the observed MEK phosphorylating activity is an indirect consequence of expression of a kinase-dead BRAF, we evaluated whether BRAF^Δ600-618^ has intrinsic kinase activity using a MEK phosphorylation assay (Bondzi et al., 2000). Briefly, cell lysates from HEK293 cells expressing the series of FLAG-tagged BRAF mutant proteins were immunoprecipitated with anti-FLAG, then precipitates were incubated with recombinant MEK1 in the presence of ATP. Phosphorylation of MEK1 was assayed by western blot against phosphorylated Ser217/221-MEK. In Figure 2, FLAG-BRAF^V600E^ immunoprecipitates had robust MEK phosphorylating activity, as expected from a class I BRAF mutant. Although less than with BRAF^V600E^, MEK phosphorylating activity was higher in immunoprecipitates of BRAF^Δ600-618^ when compared to empty vector control. This increased phosphorylation activity was similar to the level observed with BRAF^D594A^, the class III mutant which lacks intrinsic kinase activity. From this, we concluded that the complexes immunoprecipitated from FLAG-BRAF^Δ600-618^ expressing cells had MEK phosphorylating activity, however we could not determine whether this activity resided in BRAF^Δ600-618^ itself or was associated with another component of the complex.

**Figure 2.**
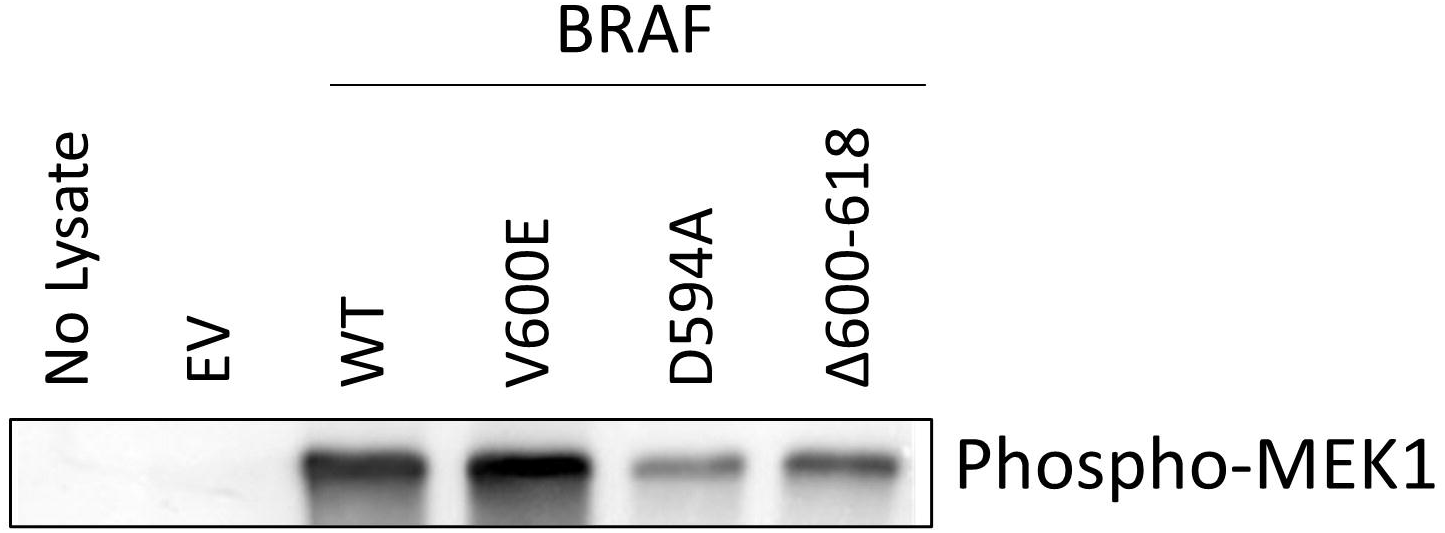
MEK phosphorylation assay of anti-FLAG immunoprecipitates from HEK293 cells ectopically expressing BRAF^Δ600-618^ and other BRAF mutants. Complexes containing FLAG-tagged BRAF proteins were immunoprecipitated from transfected HEK293 cell lysates and incubated with recombinant MEK.

### Pathway activation by BEAF^Δ600-618^ is suppressed by the introduction of a dimerization-deficient mutation

To distinguish between these scenarios, we introduced D509H mutations into each construct to disrupt BRAF dimerization (Roring et al., 2012). Previous work has shown that class I BRAF mutants signal as activated monomers and do not require dimerization to activate the MEK/ERK pathway. Class II and III BRAF mutants require dimerization to activate the MEK/ERK pathway.

Figure 3 shows MEK and ERK phosphorylation in lysates of cells expressing BRAF proteins with and without the D509H mutation. The class I mutant, BRAF^V600E^ showed similar levels of phosphorylated ERK and MEK with and without the D509H mutation indicating dimerization was not required for downstream activation of ERK and MEK. MEK and ERK activity were reduced but not eliminated when the D509H mutation was introduced into BRAF^WT^. Introduction of the D509H mutation strongly reduced MEK and ERK activity levels induced by BRAF^Δ600-618^. This indicates that BRAF^Δ600-618^ requires binding to wild-type BRAF or another RAF family member to promote pathway activation. The D509H mutation also suppressed MEK/ERK activity in BRAF^D594A^ expressing cells, an expected result because this class III BRAF mutation was previously reported to require dimerization to activate downstream signaling of MEK and ERK (Yao et al., 2015).

**Figure 3.**
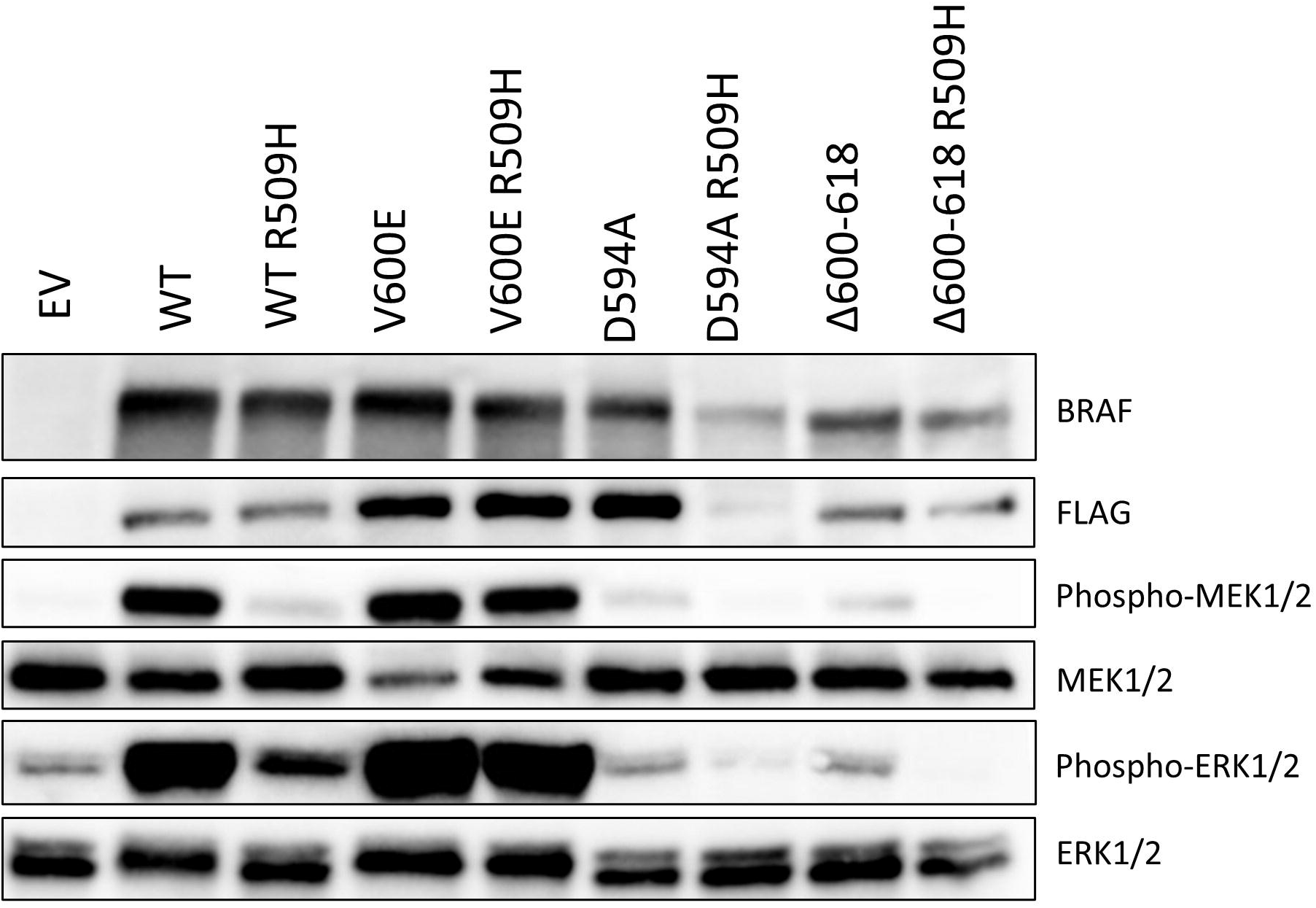
Pathway activation by BRAF^Δ600-618^ is suppressed by the introduction of a dimerization-deficient mutation. MEK/ERK activation was analyzed by western blotting of total cell lysates of transfected HEK293 cells.

### Introduction of a dimerization-deficient mutation eliminates the MEK phosphorylating activity in the BRAF^Δ600-618^ immunoprecipitates

The introduction of dimerization-deficient mutations allowed us to directly evaluate whether the kinase activities detected in the immunoprecipitates in Fig. 2 resided in the target protein or with other components of the immunoprecipitated complex. Fig. 4 shows that introduction of the D509H mutation eliminated the MEK phosphorylating activity in BRAF^Δ600-618^ immunoprecipitates. Immunoprecipitates of the known kinase-dead mutant, BRAF^D594A^, similarly lost MEK phosphorylating activity. These results suggest that, like BRAF^D594A^, BRAF^Δ600-618^ is a class III kinase-dead mutant which retains the ability to promote pathway activation by recruitment of other RAF family members.

**Figure 4.**
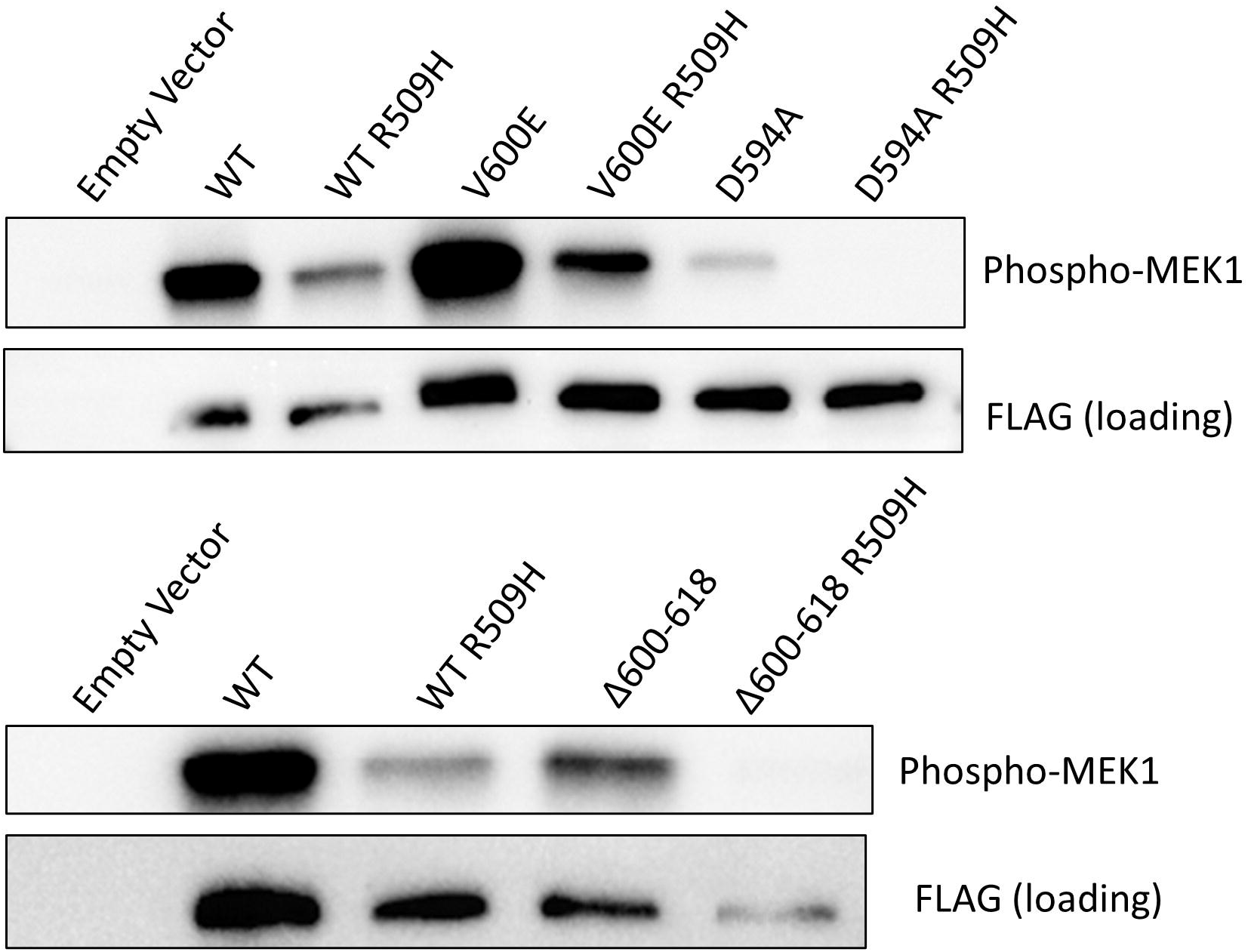
MEK phosphorylation assay of anti-FLAG immunoprecipitates from HEK293 cells ectopically expressing BRAF^Δ600-618^ and other BRAF mutants, each also containing the R509H dimerization-deficiency mutation. Introduction of the R509H mutation eliminated the MEK phosphorylating activity in the BRAF^Δ600-618^ immunoprecipitates. Anti-FLAG western blot on these membranes confirmed immunoprecipitation of FLAG-BRAF under each condition.

**Figure 5.**
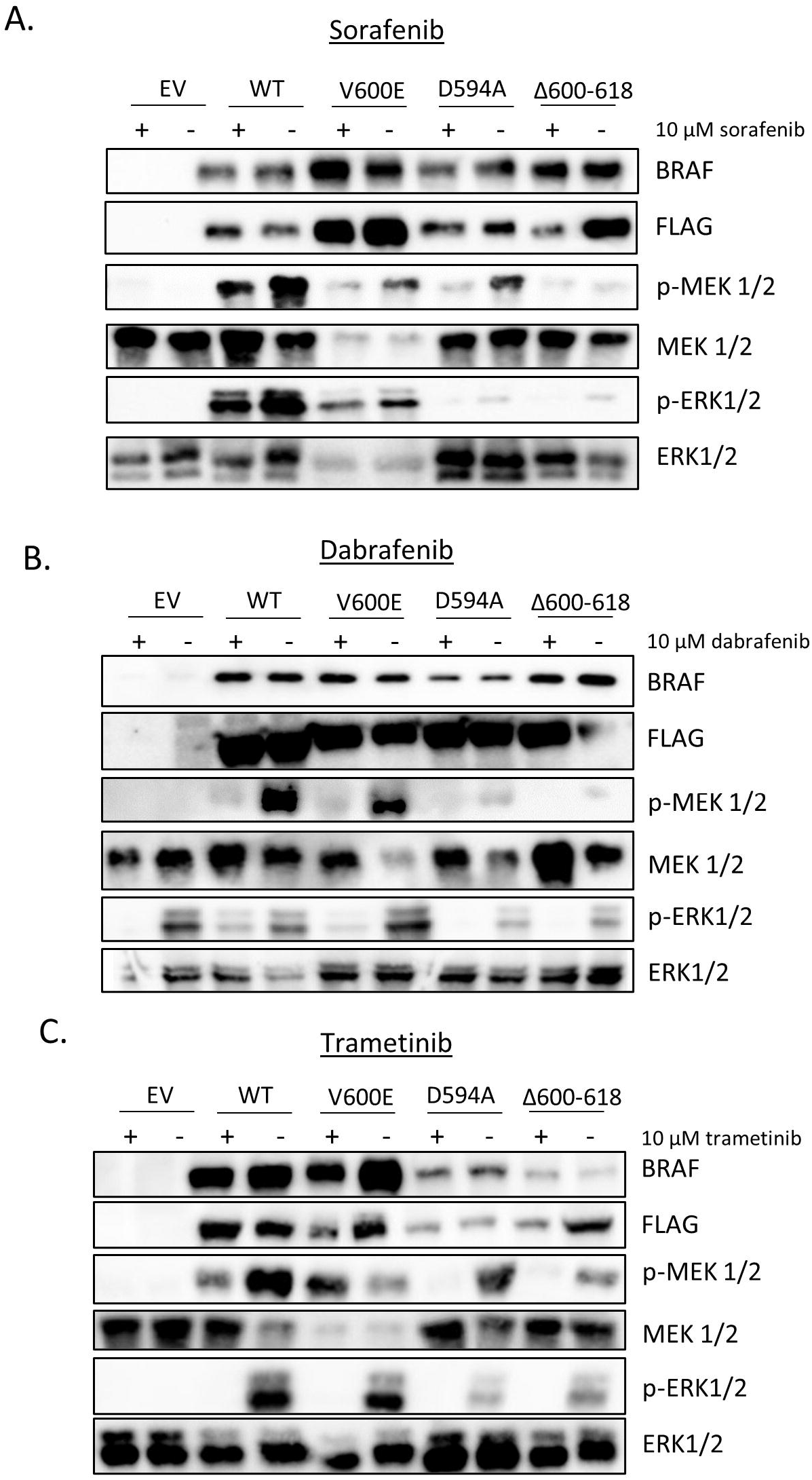
Evaluation of the sensitivity of MEK/ERK pathway activity in HEK293 cells ectopically expressing BRAF^Δ600-618^ and other BRAF mutations to (A) Sorafenib, (B) Dabrafenib and (C) Trametinib.

### Response of BRAF^Δ600-618^ expressing cells to BRAF and MEK inhibitors

While therapeutic options for several BRAF^V600E^ mutant cancers have been well established, the appropriate treatment strategy for non-classical BRAF mutations is still under investigation. We evaluated whether a pan-RAF inhibitor (sorafenib), an inhibitor of BRAF^V600E^ (dabrafenib), and a MEK inhibitor (trametinib) could reduce pathway activity elicited by the BRAF^Δ600-618^ mutant. All three inhibitors suppressed MEK/ERK activity in BRAF^Δ600-618^-expressing cells under the conditions tested. When compared with the response profiles of the other BRAF mutants analyzed, the effects observed with BRAF^Δ600-618^ were most similar to BRAF^D594A^.

## DISCUSSION

We found that ectopic expression of BRAF^Δ600-618^ mutant caused elevated MEK/ERK pathway activity, which is consistent with the hypothesis that this represents an oncogenic mutation. This pathway activation was not a direct result of BRAF^Δ600-618^-mediated MEK phosphorylation. Rather, we determined that BRAF^Δ600-618^ lacks intrinsic kinase activity. Instead, the BRAF^Δ600-618^ mutant participated in a complex which recruited the MEK phosphorylating activity of an endogenous RAF kinase family member. The oncogenic activity of BRAF^Δ600-618^ was abolished by introducing a dimerization deficiency mutation. Accordingly, we concluded that this large inframe deletion in the kinase domain met the criteria for a class III BRAF mutation and the activity of BRAF^Δ600-618^ in our assays generally paralleled that of BRAF^D594A^, a recognized member of this class (Yao et al., 2017).

More than half of all malignant melanoma patients harbor a BRAF mutation. While treatment options for the most common mutation, V600E, have been well defined, treatment for the 34% of melanoma patients harboring a non-V600 mutations is less clear (Carter et al., 2015). In particular, it may be challenging to infer solely from sequence data whether a novel non-V600 mutant may fit in either Class II or III or, alternatively, may be simply an irrelevant passenger mutation. The ability to characterize these non-V600 mutations may assist with appropriate treatment selection for patients.

In this instance, short-term treatment with a BRAF inhibitor (dabrafenib), a pan-RAF inhibitor (sorafenib) and a MEK inhibitor (trametinib) all caused diminished pathway activity in cells ectopically expressing BRAF^Δ600-618^. With the relatively high concentrations of inhibitors used, this experiment is probably most usefully interpreted as providing insight into the factors on which pathway activation by this unusual BRAF mutation depend, rather than providing actionable guidance on therapy choice in this setting. Consistent with other class III mutations, these data indicate that RAF and MEK activity remain important. Key factors relevant to therapeutic response that are not captured in this experiment are the duration of pathway suppression that can be achieved, as well as the influence of the likely obligate RAS activation that co-occurs with these BRAF mutations in tumors and of any signaling feedback that may occur over longer periods in that context.

Clinical experience of using single agent BRAF (Mazieres et al., 2020) or MEK (Johnson et al., 2020) inhibitors has not been positive, and this particular patient did not respond to dual dabrafenib/trametinib therapy. The patient in whom we identified this mutation had only single gene BRAF testing, so whether co-occurring mutations in RAS or other genes might have influenced the lack of response to dabrafenib/trametinib is not clear. RAF dimer inhibitors such as LY3009120 (Sullivan et al., 2020) and BGB-283 (Desai et al., 2020) or ERK inhibitors such as ulixertinib (Sullivan et al., 2018) may prove to be more promising for this class of mutations in future.

## Acknowledgements

This work was supported by the Gundersen Medical Foundation. We thank Mark Burkard, MD, PhD for useful discussions. PAK holds the Dr. Jon & Betty Kabara Endowed Chair in Precision Oncology. MAG was supported by a Norman L. Gillette Jr Postdoctoral Fellowship. MEC was supported by a Kabara – St. Mary’s University of Minnesota Summer Fellowship.

## REFERENCES

Barzon, L., Masi, G., Boschin, I.M., Lavezzo, E., Pacenti, M., Casal Ide, E., Toniato, A., Toppo, S., Palu, G., and Pelizzo, M.R. (2008). Characterization of a novel complex BRAF mutation in a follicular variant papillary thyroid carcinoma. Eur J Endocrinol 159, 77–80.

Bondzi, C., Grant, S., and Krystal, G.W. (2000). A novel assay for the measurement of Raf-1 kinase activity. Oncogene 19, 5030–5033.

Burkard, M.E., Deming, D.A., Parsons, B.M., Kenny, P.A., Schuh, M.R., Leal, T., Uboha, N., Lang, J.M., Thompson, M.A., and Warren, R. (2017). Implementation and clinical utility of an integrated academic-community regional molecular tumor board. JCO Precision Oncology 1, 1–10.

Carter, J., Tseng, L.H., Zheng, G., Dudley, J., Illei, P., Gocke, C.D., Eshleman, J.R., and Lin, M.T. (2015). Non-p.V600E BRAF Mutations Are Common Using a More Sensitive and Broad Detection Tool. Am J Clin Pathol 144, 620–628.

Davies, H., Bignell, G.R., Cox, C., Stephens, P., Edkins, S., Clegg, S., Teague, J., Woffendin, H., Garnett, M.J., Bottomley, W., et al. (2002). Mutations of the BRAF gene in human cancer. Nature 417, 949–954.

Desai, J., Gan, H., Barrow, C., Jameson, M., Atkinson, V., Haydon, A., Millward, M., Begbie, S., Brown, M., Markman, B., et al. (2020). Phase I, Open-Label, Dose-Escalation/Dose-Expansion Study of Lifirafenib (BGB-283), an RAF Family Kinase Inhibitor, in Patients With Solid Tumors. J Clin Oncol 38, 2140–2150.

Flaherty, K.T., Infante, J.R., Daud, A., Gonzalez, R., Kefford, R.F., Sosman, J., Hamid, O., Schuchter, L., Cebon, J., Ibrahim, N., et al. (2012). Combined BRAF and MEK inhibition in melanoma with BRAF V600 mutations. N Engl J Med 367, 1694–1703.

Greaves, W.O., Verma, S., Patel, K.P., Davies, M.A., Barkoh, B.A., Galbincea, J.M., Yao, H., Lazar, A.J., Aldape, K.D., Medeiros, L.J., et al. (2013). Frequency and spectrum of BRAF mutations in a retrospective, single-institution study of 1112 cases of melanoma. J Mol Diagn 15, 220–226.

Heidorn, S.J., Milagre, C., Whittaker, S., Nourry, A., Niculescu-Duvas, I., Dhomen, N., Hussain, J., Reis-Filho, J.S., Springer, C.J., Pritchard, C., et al. (2010). Kinase-dead BRAF and oncogenic RAS cooperate to drive tumor progression through CRAF. Cell 140, 209–221.

Johnson, D.B., Zhao, F., Noel, M., Riely, G.J., Mitchell, E.P., Wright, J.J., Chen, H.X., Gray, R.J., Li, S., McShane, L.M., et al. (2020). Trametinib Activity in Patients with Solid Tumors and Lymphomas Harboring BRAF Non-V600 Mutations or Fusions: Results from NCI-MATCH (EAY131). Clin Cancer Res 26, 1812–1819.

Mazieres, J., Cropet, C., Montane, L., Barlesi, F., Souquet, P.J., Quantin, X., Dubos-Arvis, C., Otto, J., Favier, L., Avrillon, V., et al. (2020). Vemurafenib in non-small-cell lung cancer patients with BRAF(V600) and BRAF(nonV600) mutations. Ann Oncol 31, 289–294.

Rogiers, A., Vander Borght, S., Tuand, K., Wolter, P., Stas, M., Boecxstaens, V., Garmyn, M., van den Oord, J.J., Vandenberghe, P., and Bechter, O. (2017). Response to targeted therapy in two patients with metastatic melanoma carrying rare BRAF exon 15 mutations: A598_T599insV and V600_K601delinsE. Melanoma Res 27, 507–510.

Roring, M., Herr, R., Fiala, G.J., Heilmann, K., Braun, S., Eisenhardt, A.E., Halbach, S., Capper, D., von Deimling, A., Schamel, W.W., et al. (2012). Distinct requirement for an intact dimer interface in wild-type, V600E and kinase-dead B-Raf signalling. EMBO J 31, 2629–2647.

Sullivan, R.J., Hollebecque, A., Flaherty, K.T., Shapiro, G.I., Rodon Ahnert, J., Millward, M.J., Zhang, W., Gao, L., Sykes, A., Willard, M.D., et al. (2020). A Phase I Study of LY3009120, a Pan-RAF Inhibitor, in Patients with Advanced or Metastatic Cancer. Mol Cancer Ther 19, 460–467.

Sullivan, R.J., Infante, J.R., Janku, F., Wong, D.J.L., Sosman, J.A., Keedy, V., Patel, M.R., Shapiro, G.I., Mier, J.W., Tolcher, A.W., et al. (2018). First-in-Class ERK1/2 Inhibitor Ulixertinib (BVD-523) in Patients with MAPK Mutant Advanced Solid Tumors: Results of a Phase I Dose-Escalation and Expansion Study. Cancer Discov 8, 184–195.

Wan, P.T., Garnett, M.J., Roe, S.M., Lee, S., Niculescu-Duvaz, D., Good, V.M., Jones, C.M., Marshall, C.J., Springer, C.J., Barford, D., et al. (2004). Mechanism of activation of the RAF-ERK signaling pathway by oncogenic mutations of B-RAF. Cell 116, 855–867.

Yao, Z., Torres, N.M., Tao, A., Gao, Y., Luo, L., Li, Q., de Stanchina, E., Abdel-Wahab, O., Solit, D.B., Poulikakos, P.I., et al. (2015). BRAF Mutants Evade ERK-Dependent Feedback by Different Mechanisms that Determine Their Sensitivity to Pharmacologic Inhibition. Cancer Cell 28, 370–383.

Yao, Z., Yaeger, R., Rodrik-Outmezguine, V.S., Tao, A., Torres, N.M., Chang, M.T., Drosten, M., Zhao, H., Cecchi, F., Hembrough, T., et al. (2017). Tumours with class 3 BRAF mutants are sensitive to the inhibition of activated RAS. Nature 548, 234–238.

Zaman, A., Wu, W., and Bivona, T.G. (2019). Targeting Oncogenic BRAF: Past, Present, and Future. Cancers (Basel) 11.

